# Coherence-based interhemispheric EEG functional connectivity changes in distinct frequency bands during eyes open meditation

**DOI:** 10.1101/2023.02.06.527252

**Authors:** G Pradeep Kumar, Kanishka Sharma, A Adarsh, Amrutha Manvi, G Ramajayam, A G Ramakrishnan

## Abstract

Meditation is a self-regulatory process practiced primarily to reduce stress, manage emotions and mental health. The objective was to study the information exchange between symmetric electrodes across the hemispheres during meditation using functional connectivity (FC) measures. We investigate the changes in the coherence between EEG electrode pairs during the meditation practiced by long-term Brahmakumaris Rajyoga meditators with open eyes and during listening to music by controls as the comparable task. Two distinct FC measures derived from coherency, namely, magnitude squared coherence (MSC) and imaginary part of coherency (ICoh) are used to study the changes in interhemispheric coherence. During baseline conditions, higher MSC is found in meditators in frontocentral and centroparietal regions and higher |ICoh| globally in higher beta and gamma bands than controls. Further, in meditators, the MSC significantly increases in higher theta and alpha bands in the frontal and parietal regions and |ICoh| significantly decreases across all regions and bands except in the alpha band during meditation. However, the control subjects with no knowledge of meditation show no change in theta or alpha MSC or |ICoh| during the music session. Distinct patterns of changes are observed with the two FC measures in different frequency bands during meditation in the meditators and music-listening session in the control subjects indicating varied information processing between the right and left hemispheres and differences in the FC measures used. We found increased MSC and decreased |ICoh| between the frontal electrodes implying increased self-awareness in meditators. The MSC between the occipital electrodes in meditators is less than the controls in baseline condition indicating a possible modified visual information processing in Rajyoga meditators due to the long-term practice of meditation with open eyes. Overall, the changes in MSC indicate increased functional integration during meditation supporting the hypothesis of cortical integration theory.

## Introduction

Meditation is a mental exercise wherein an individual focuses attention on a single object, thought, experience, or concept in order to improve the core psychological, emotional, and attention regulation (Travis, 2014). Meditation may involve distinct complex practices based on different techniques and philosophies (Tang et al., 2015; Wallace and Benson, 1972; West, 1979). Brain connectivity relates to anatomical or functional connectivity (FC) across the brain regions. Neuronal oscillations and their synchronicity resulting in functional networking across different brain regions can be measured using electroencephalography (EEG) (Bowyer, 2016).

EEG coherence is an FC measure that quantifies the brain’s network formation involving highly shared activity and integration (Nunez, 2000). The nature of this frequency-correlated activity varies with the cognitive state. Coherence is considered to be the second marker of frontal lobe functioning. A high value of EEG coherence and synchrony is associated with functional connection, integration (Hebert et al., 2005) and information exchange (Petsche et al., 1997; Pfurtscheller and Andrew, 1999) between the corresponding brain regions. Alpha coherence has been found to be an important predictor of executive function. A low level of EEG coherence is associated with normal aging (Koyama et al., 1997), schizophrenia (Wada et al., 1998), decreased cerebral blood flow, large white matter lesions, and poor outcome of treatment for depression (Leuchter et al., 1997). Knott et al. (Knott et al., 2001) reported reduced interhemispheric coherence (IHC) in several cortical areas in each of the EEG bands in depressive male patients. Interhemispheric magnitude squared coherence (MSC) has been used as a metric to differentiate Alzheimer’s disease patients from healthy elderly subjects (Sankari et al., 2011). Liston et al (Liston et al., 2009) found reduced attentional control and disrupted FC in the prefrontal cortex of healthy adults during periods of psychosocial stress. A decrease in IHC with age indicates a reduction in interhemispheric functional coupling for both females and males. The decrease in EEG synchronization is postulated to be due to the alterations in the corpus callosum and the white matter, reducing the connectivity (Duffy et al., 1996).

IHC has been suggested as a potential marker in many clinical studies related to mental health. Compared to healthy controls, lower inter-hemispheric delta coherence has been observed in frontal and parietal regions in anxiety, depression, and ADHD patients, and increased frontal and central coherence of theta in EO condition in depression. Central theta coherence tends to be high in patients with anxiety, depression, ADHD, and Asperger’s syndrome (Markovska-Simoska et al., 2018). In age-related disorders such as dementia and mild cognitive impairment, alpha and theta coherence decrease as the manifestation of impairment in connectivity between frontal executive centers and the parietal and temporal processing areas. Also, the long-range network connectivity is compromised (Babiloni et al., 2016). In naive schizophrenic patients, the central region had lower IHC in both eyes closed and eyes open conditions, and the parietal region in eyes closed condition (John et al., 2002). A study on healthcare professionals during COVID-19 reported decreased IHC in theta and alpha bands in the parietal region indicating early signs of negative impact due to stress exposure (LoMauro et al., 2022). Reduction of interhemispheric connectivity in the prefrontal cortex was observed during stress as compared to control situation (Al-Shargie, 2019).

Various neurophysiological studies have been conducted on the changes in the conscious state during meditation using parameters such as EEG interhemispheric coherence. However, less is known about the involvement of coordinated communication for effective information processing between brain regions across the two hemispheres. Higher levels of alpha EEG coherence during transcendental meditation (TM) are correlated to facilitated cognition and heightened awareness. An increase in alpha coherence from the frontal to central regions of the brain was reported during TM and it was suggested that coherence was a more sensitive parameter than power or amplitude changes in EEG (Dillbeck and Bronson, 1981; Travis and Wallace, 1999). An increase in slow alpha interhemispheric EEG coherence was reported in the frontal region of the brain during Zen meditation (Murata et al., 2004). However, this study used only two electrode pairs in the frontal and central lobes. A study involving EEG data from F3 and F4 electrodes reported decreased coherence during individual meditation and increased coherence during a group meditation, both in the alpha band in TM and Zen meditators (Newandee et al., 1996). Higher frontal coherence and lateral asymmetry were observed in TM subjects than in controls (Travis and Arenander, 2006), and coherence was reported to be more sensitive than lateral asymmetry.

Another study on five different meditations using lagged coherence reported a predominant decrease in coherence during meditation in all five traditions across all the eight frequency bands considered (Lehmann et al., 2012). Meditators practicing mindfulness, Zen, and Vipassana have shown higher levels of focus and decreased mind-wandering in EEG during the practice (Rodriguez-Larios et al., 2021). In another study, experienced meditators had higher maximum betweenness centrality than novice meditators in alpha band EEG indicating increased integration of the networks which may be utilized for studying the quality of meditation (van Lutterveld et al., 2017). Increased information processing has been postulated with enhanced microstructural properties of interhemispheric white matter network of the corpus callosum in long-term Rajyoga meditators (Sharma et al., 2018b). Highly creative individuals maintain a higher alpha rhythm during the act of creating than during rest with the eyes closed (Martinda and Hines, 1973).

Various studies have been conducted on the effect of Rajyoga meditation based on physiological, neurophysiological, and psychological measures. Increased grey matter volume was observed in reward processing centres associated with happiness in Rajyoga meditators compared to non-meditators. Happiness score was higher in meditators than controls and was positively correlated with experience (Babu et al., 2020). A study reported a significant reduction in anxiety, depression, obsessive-compulsive disorder, and chronic tension headache in the meditation intervention group compared to the control group (Arora et al., 2014), (Amritsar, 2014), (Mehta et al., 2020). A similar study on patients undergoing cardiac surgery reported reduced anxiety and cortisol levels in meditators compared to the control group (Kiran et al., 2017) and improved cardiorespiratory functions due to long-term practice of Rajyoga meditation (Sukhsohale and Phatak, 2012). Rajyoga meditators reported increased happiness, self-satisfaction scores (Ramesh et al., 2013), and only 4.7% of the 801 meditators in a study reported moderate and severe psychological outbreaks due to Covid 19 (Madhu et al., 2022). Reduced delta and increased low-alpha activity during Rajyoga meditation was reported in a localization study using EEG (Sharma et al., 2020). Long-term Rajyoga meditators could change their mental states rapidly after an attentional task indicating the neuroplastic changes in facilitating quick shifts as compared to short-term meditators and non-meditators (Nair et al., 2017). A study on changes in default mode network (DMN) in Rajyoga meditators during meditation revealed that the frequency and duration of the DMN microstates were higher than healthy controls indicating the trait effects due to long-term meditation practice (Panda et al., 2016). The decreased distance between covariance matrices obtained from successive epochs was reported in the frontal lobe indicating increased uniformity during Rajyoga meditation (Ganesan et al., 2020). Entropy in long-term Rajyoga meditators was higher during meditation than the baseline condition which is an indication of a non-pharmacologically induced state of high entropy (Kumar et al., 2021).

Integration is required to enable information processing across distributed brain regions to execute higher-level processes for cognition. Long-range interactions in different frequency bands are involved during various cognitive functions. This has been postulated as the cortical integration theory by (Hebert et al., 2005), (Von Stein and Sarnthein, 2000). There has been no reported, systematic study on changes in interhemispheric EEG spectral coherence during modified states of consciousness. Most studies have emphasized changes in FC across all the electrode pairs. In a previous study by sharma et. al (Sharma et al., 2018a) alpha asymmetry across hemispheres was found to be higher in long-term Rajyoga meditators in frontal and parietal regions. In continuation with the study on Rajyoga meditation, our objective was to examine information exchange across the hemispheres between symmetric electrodes and its implications for functional integration. The results reported are confined to interhemispheric FC between symmetric electrodes. FC between anterior and posterior regions of the brain, and intrahemispheric FC shall be analyzed later. The present study examines the interhemispheric functional connectivity using measures based on coherency during the modified state of consciousness due to Brahmakumaris Rajyoga (BKRY) meditation in long-term meditators, listening to music in controls and during task-free resting before and after meditation or music sessions.

## Results

Two types of analyses are performed, the first to look for the state changes that occur in the baseline conditions before and after performing meditation (MED) in meditators and due to music (MUS) session in controls. The second analysis is to look for the trait changes by comparing the difference in baseline FC measures between the control subjects and the meditators. Only the electrode pairs showing a magnitude ΔMSC ≥ 0.03 are considered for the statistical significance test due to the global changes in MSC and sensitivity to referencing. No such threshold was applied for |ICoh| analysis.

### Changes across conditions in meditators

Figures 3a and 3b give the plots of change in MSC values (ΔMSC) due to the task in meditators and control subjects, respectively, as a function of frequency. The vertical dashed lines represent the boundaries between the different frequency bands. The results reveal an increase in MSC during meditation (MED) compared to the baseline eyes-open (EO1_M_) condition. Frontal, frontocentral, and centroparietal lobes show an increase in MSC during meditation in the delta and alpha bands. The O1-O2 electrode pair in the occipital lobe is prominent in MSC in all the bands except alpha. However, the coherence in the central lobe decreases during MED in the slow gamma band which needs further exploration (Fig. 3a). Similar to the increase during MED over the EO1_M_ condition, MSC values are higher during MED than in the post-meditation eyes-open (EO2_M_) condition. The number of statistically significant pairs reduces drastically across all the bands which may be the aftereffect of meditation (Supplementary Fig. 8a). Specifically, the number of electrode pairs that have increased coherence is reduced during MED when compared with EO2_M_ condition. Compared to the initial EO1_M_ condition (Supplementary Fig. 8d), MSC in lower frequency bands is higher in the EO2_M_ baseline mostly in electrode pairs in frontal and frontocentral regions. MSC differences were present between eyes closed conditions (EC1_M_, EC2_M_) and MED, mostly in central and parietal regions (Supplementary Fig. 8b and Fig. 8c. However, there are many statistically significant pairs between EO1_M_, EO2_M_ and eyes closed conditions in both pre and post-meditation sessions (EC1_M_, EC2_M_) in posterior region in alpha, beta, and slow gamma bands (Supplementary Fig. 8e, Fig. 8f, Fig. 8h and Fig. 8i). Compared with the EC1_M_ condition (Supplementary Fig. 8g), changes in MSC values are seen in EC2_M_ condition. An increase is observed in the anterior part of the cortex in the theta and alpha bands and in the centroparietal region in the beta band. A decrease in MSC is observed in the parieto-occipital and occipital regions in beta and slow gamma bands.

**Figure 1:**
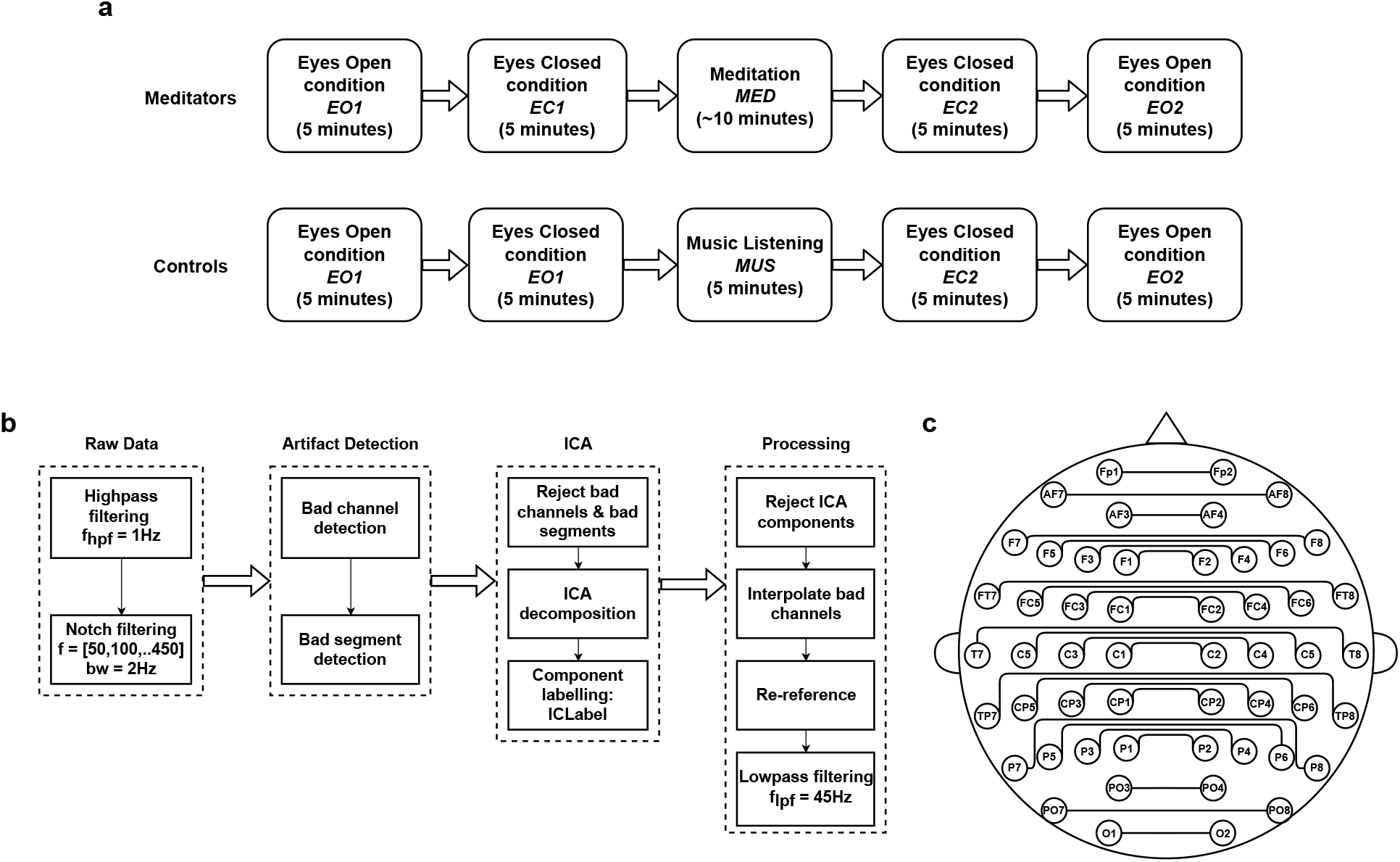
**a)** The recording protocol used for EEG data collection from **(top:)** meditators performing Rajyoga meditation and **(bottom:)** control subjects listening to music. **b)** the preprocessing pipeline. **c)** Interhemispheric electrode pairs considered for the analysis of change in MSC and |ICoh| in the study.

**Figure 2:**
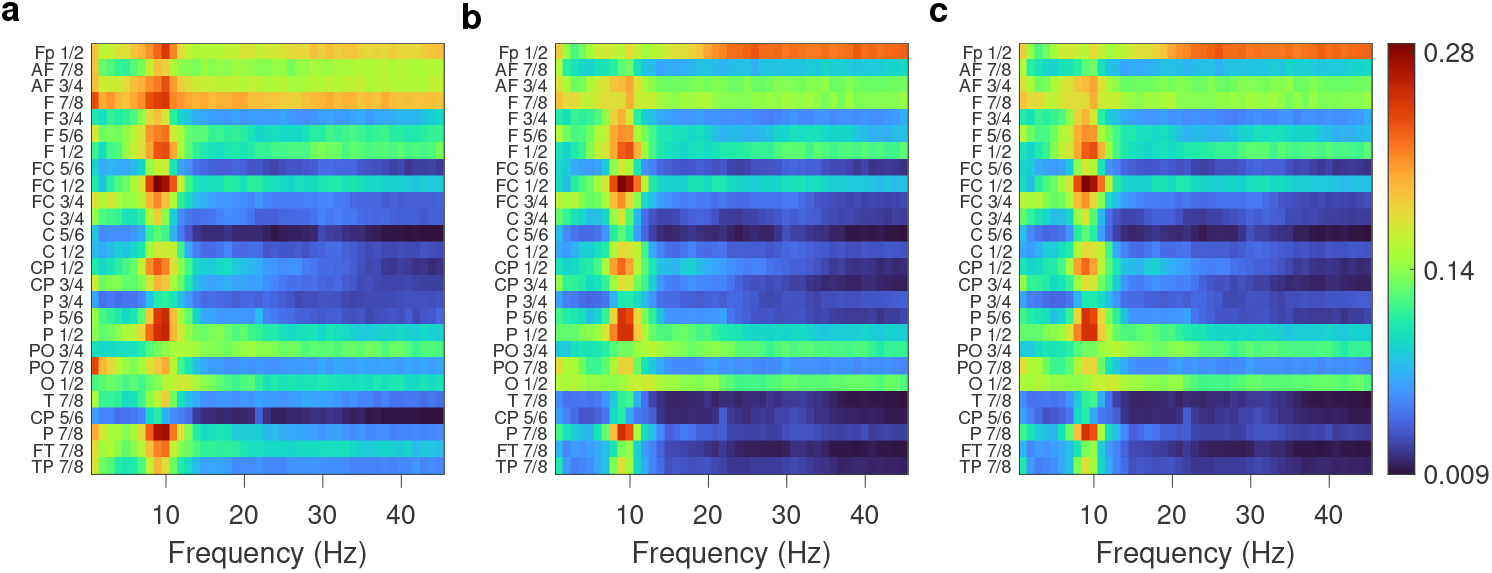
Plots of MSC values between symmetric electrode pairs versus frequency in meditators during meditation with different referencing schemes: **a)** average mastoids **b)** common average **c)** REST reference. All the colormaps have been plotted using the same scale.

**Figure 3:**
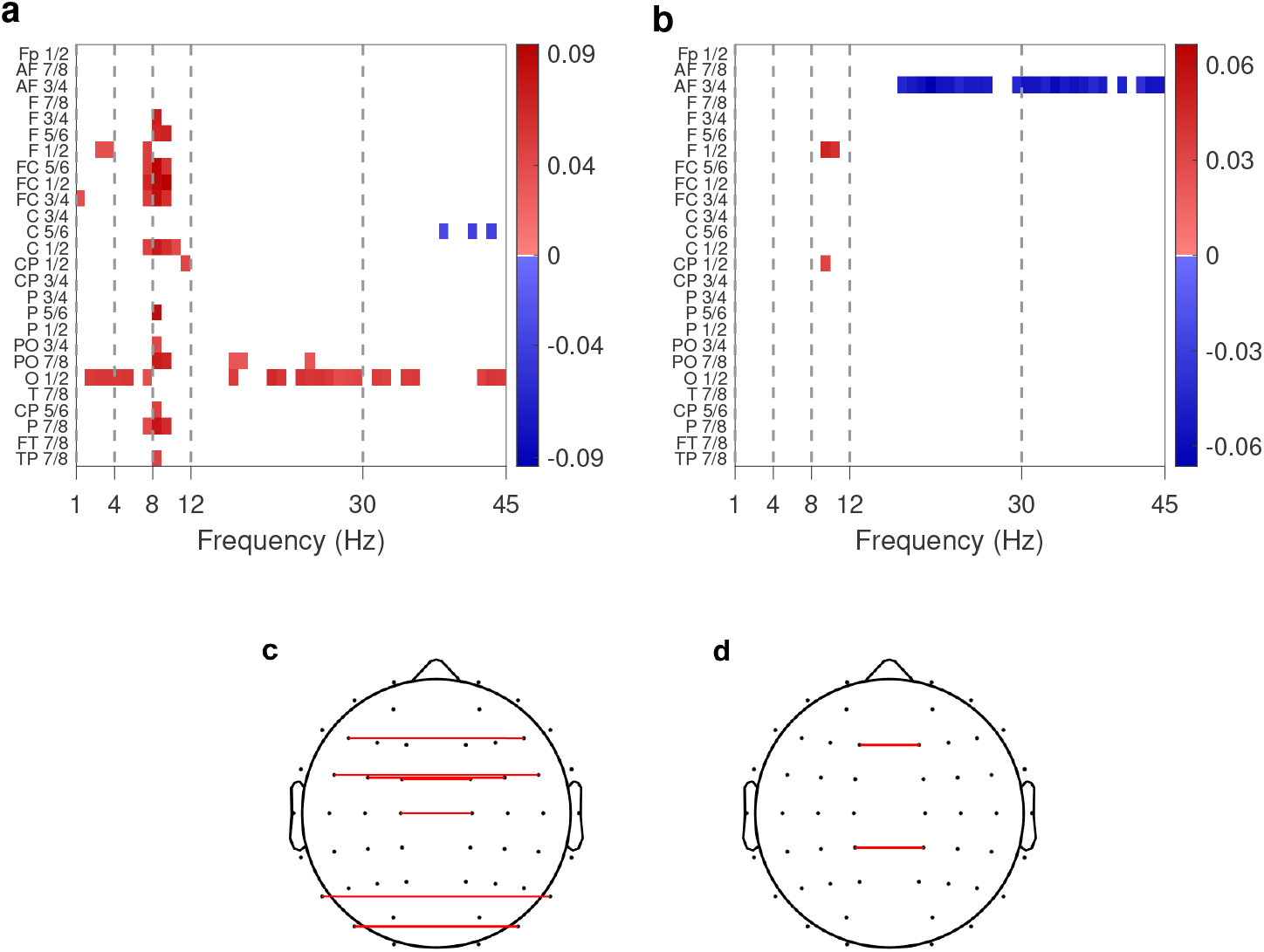
Plots of ΔMSC between symmetric electrode pairs due to task versus frequency. **a)** MED - EO1_M_ in meditators; **b)** MUS - EO1_C_ in controls. Positive (negative) ΔMSC indicates higher (lower) MSC in the first of the compared states than the second one and is represented by red (blue) color. In y-axis, from the top to the bottom, the electrode pairs are arranged according to anterior to posterior regions of the brain, with the temporal electrodes at the bottom. Only the electrode pairs and frequencies showing a ΔMSC of magnitude ≥ 0.03 and statistically significant with p ≤ 0.05 are highlighted and others are shown as zero by white color in the figure. The vertical dashed lines are the boundaries between the various frequency bands. **c)** Connected topoplots of the difference in ΔMSC between conditions within the meditators group at 9 Hz. **d)** Same as in **(c)** but within the control group. The red color indicates that the ΔMSC is higher during task conditions and blue, is the inverse.

**Figure 4:**
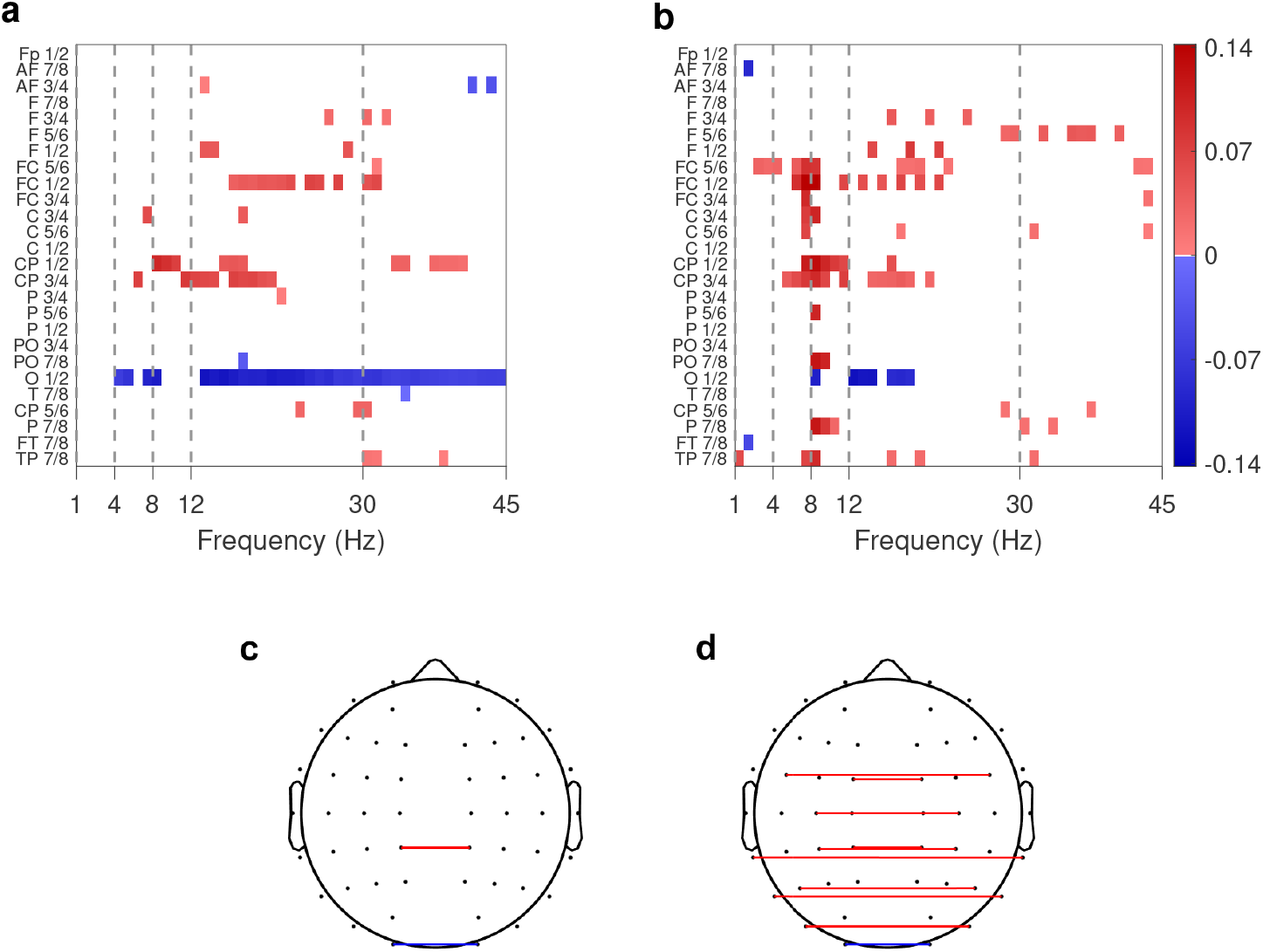
Plots of differences between the MSC (between symmetric electrode pairs) values across different groups (meditators and controls) as a function of frequency. The difference in MSC **a)** between the initial eyes-open conditions of meditators and controls, EO1_M_-EO1_C_ ; **b)** between MED and MUS sessions, MED-MUS. All the figures are plotted with the same color scale. In the y-axis, from the top to the bottom, the electrode pairs are arranged according to anterior to posterior regions of the brain, with the temporal electrodes at the bottom. Only the electrode pairs and frequencies statistically significant with p ≤ 0.05 are highlighted. **c)** Connected topoplots of the difference in MSC between baseline conditions across groups at 9 Hz. **d)** Same as in **(c)** but during the meditation and music session. The red color indicates that the MSC in meditators is higher than that of controls and blue, is the inverse. Difference values that are not significant are shown as zero with white color.

**Figure 5:**
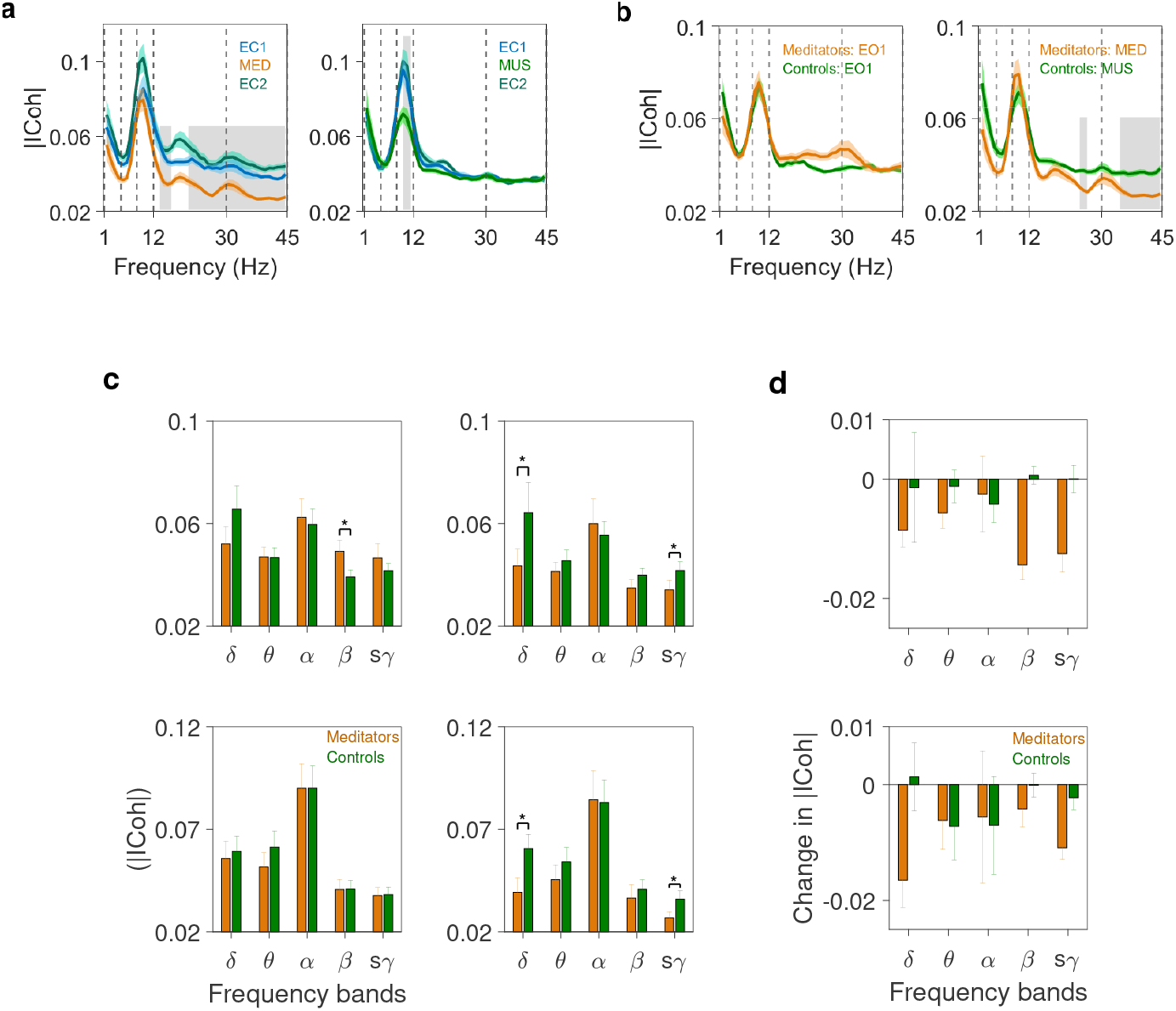
|ICoh| in meditators and controls: **a)** Grand average of |ICoh| across all the electrode pairs as a function of frequency during different conditions within the group. The respective light color band associated with each solid line represents SEM. Left column: meditators and right column: controls. The vertical dashed lines are the boundaries between the EEG freq. bands. **b)** Same as in **(a)** but across groups during EO1 (left) and MED/MUS (right) sessions. The grey shaded area indicates the region of statistical differences between the conditions (EO1&MED, EO1&MUS) and groups after Bonferroni correction for multiple comparisons. **c)** Mean |ICoh| in frontal (top) and occipital (bottom) regions during EO1 (left column) and during meditation or music sessions (right column) in different frequency bands. Error bars represent SEM. * represent the statistical significance after Bonferroni correction. **d)** Change in |ICoh| in frontal (top) and occipital (bottom) regions during MED/MUS sessions compared to EO1 (MED-EO1_M_ and MUS-EO1_C_) in different frequency bands.

**Figure 6:**
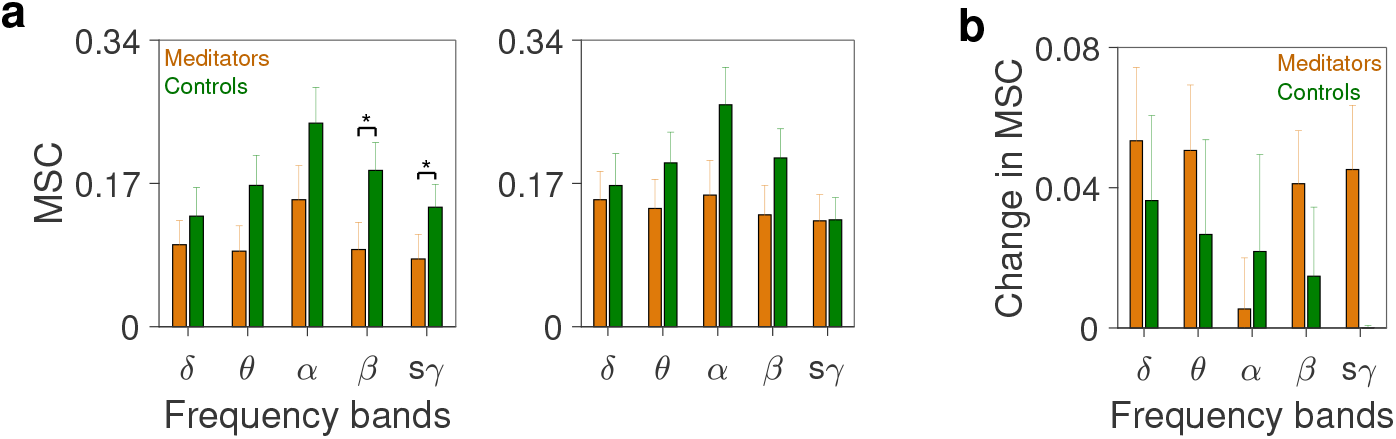
MSC between the occipital O1-O2 electrodes in meditators and controls. **a)** Mean MSC in each band during baseline eyes-open condition -EO1 (left) and during task condition (right). **b)** Change in MSC during meditation or music session with respect to EO1. Error bars indicate SEM. * represent the statistical significance after Bonferroni correction.

**Figure 7:**
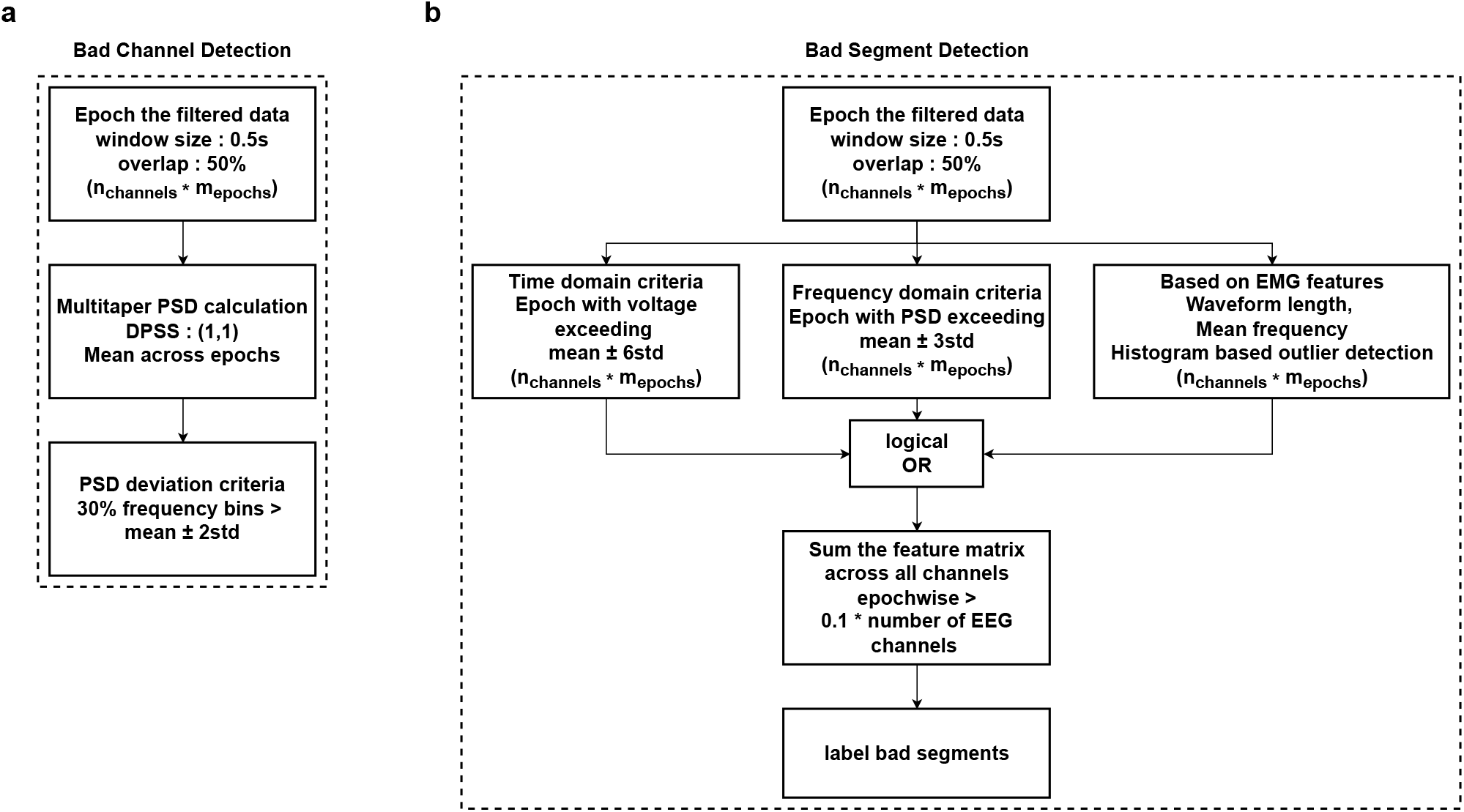
Bad channel and segment detection: **a)** Flow chart describing the steps involved in detecting bad channels with windowing, PSD calculation of the epochs, and detection criteria based on PSD. **b)** Flow chart describing the steps involved in detecting bad segments with windowing, extracting features in time, frequency domain, and based on EMG features applied to EEG channels, then bad segment detection criteria.

**Figure 8:**
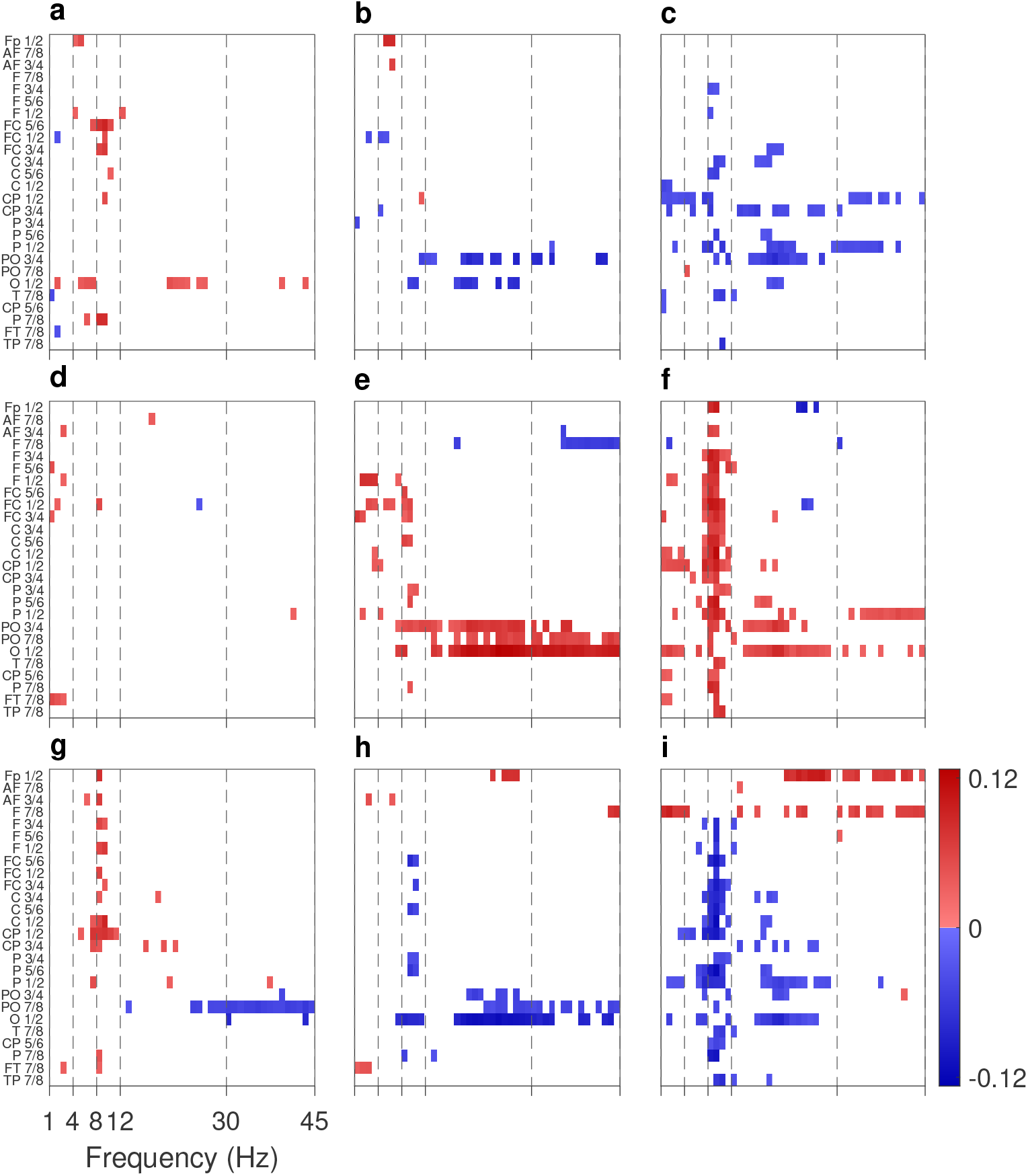
Plots of ΔMSC between symmetric electrode pairs versus frequency in meditators. **a)** MED-EO2_M_; **b)** MED-EC1_M_; **c)** MED-EC2_M_; **d)**EO2_M_-EO1_M_; **e)** EC1_M_-EO1_M_; **f)** EC2_M_-EO1_M_; **g)** EC2_M_-EC1_M_; **h)** EO2_M_-EC1_M_; **i)** EO2_M_-EC2_M_. All the other notations, etc. are exactly the same as in Fig. 2.

To identify the frequency and lobe-specific networks in experienced meditators during the baseline and meditation conditions, frequency plots of |ICoh| are obtained. The left column of Fig. 5a presents the grand average |ICoh| values of all the interhemispheric electrode pairs in EO1_M_ baseline, MED, and EO2_M_ conditions. Solid lines represent the grand average and the respective light-colored bands indicate the SEM. The plots demonstrate that in all these conditions connectivity is most prominent in delta, alpha, beta, and slow gamma bands. The results show that |ICoh| decreases significantly (at Bonferroni significance level of 0.05/45 = 0.001) during MED from the EO1_M_ values in the low frequencies (2 and 5 Hz) and also the high-frequency band from 21 to 45 Hz. The results were also significant between MED and EO2_M_ at low frequencies (2, 3, and 5 Hz) and the high-frequency band from 15 to 45 Hz. |ICoh| showed a significant decrease during MED compared to both the eyes closed EC1_M_ and EC2_M_ as given in the left column of Supplementary Fig. 11a.

### Changes across conditions in controls

In control subjects, there are negligible IHC changes in the lower frequency bands. In the higher frequency beta and slow gamma bands, MSC decreases in the anterior-frontal region of the brain during the music (MUS) session as compared to the initial eyes-open baseline (EO1_C_) condition (Fig. 3b). The coherence decreased further in these bands in frontal regions post-eyes-open (EO2_C_) condition compared to MUS session (Supplementary Fig. 9b). No clusters were found in EO2_C_ condition compared to the MUS session (Supplementary Fig. 9a). The results reveal that the MSC in the alpha band is higher in eyes closed conditions in both EC1_C_ (Supplementary Fig. 9b) and EC2_C_ session compared to MUS session except in frontal electrodes F7-F8, where the MSC increases during MUS session in higher frequency bands (Supplementary Fig. 9c). A decrease in MSC is observed in AF3-AF4 EO2_C_ condition (Supplementary Fig. 9d). MSC decreases across a pair of frontal electrodes in higher frequencies and increases across electrodes in frontocentral, central, and centroparietal regions in lower frequencies in EO2_C_ condition compared with EO1_C_ condition (Supplementary Fig. 9d). Negligible changes in IHC were observed between EO2_C_ and EO1_C_ (Supplementary Fig. 9g). The MSC differences between EO1_C_, EO2_C_ and EC1_C_, EC2_C_ conditions were mostly in the alpha band across all the regions and frontal region in higher frequency bands.

Closed conditions showed increased MSC in the alpha band and decreased MSC in frontal regions in higher frequency bands (Supplementary Fig. 9e, Fig. 9f and Fig. 9i). It is clear from Figs. 3a, 3b, and Supplementary Figs. 8, 9 that the number of electrode pairs showing significant changes in IHC during the task is higher in meditators than controls. However, the pattern of IHC across different conditions differs in meditators from the controls. The frequency plots of |ICoh| given in Fig. 5a right column, reveal no significant change in the grand average of |ICoh| values across EO1_C_ and EO2_C_ conditions compared with a music session in any band except the slow gamma. A decrease in |ICoh| values during MUS session was observed in the frequency band 30 - 37 Hz for p ≤ 0.05 which is not significant at Bonferroni significant levels supporting the results of the MSC given in Fig. 3b. The frequency plots of |ICoh| given in the right column of Fig. 11a, showed no significant change in the grand average of |ICoh| values across EC1_C_ and EC2_C_ conditions compared with a music session in any band except in alpha, which may be due to the eyes open condition during MUS session.

**Figure 9:**
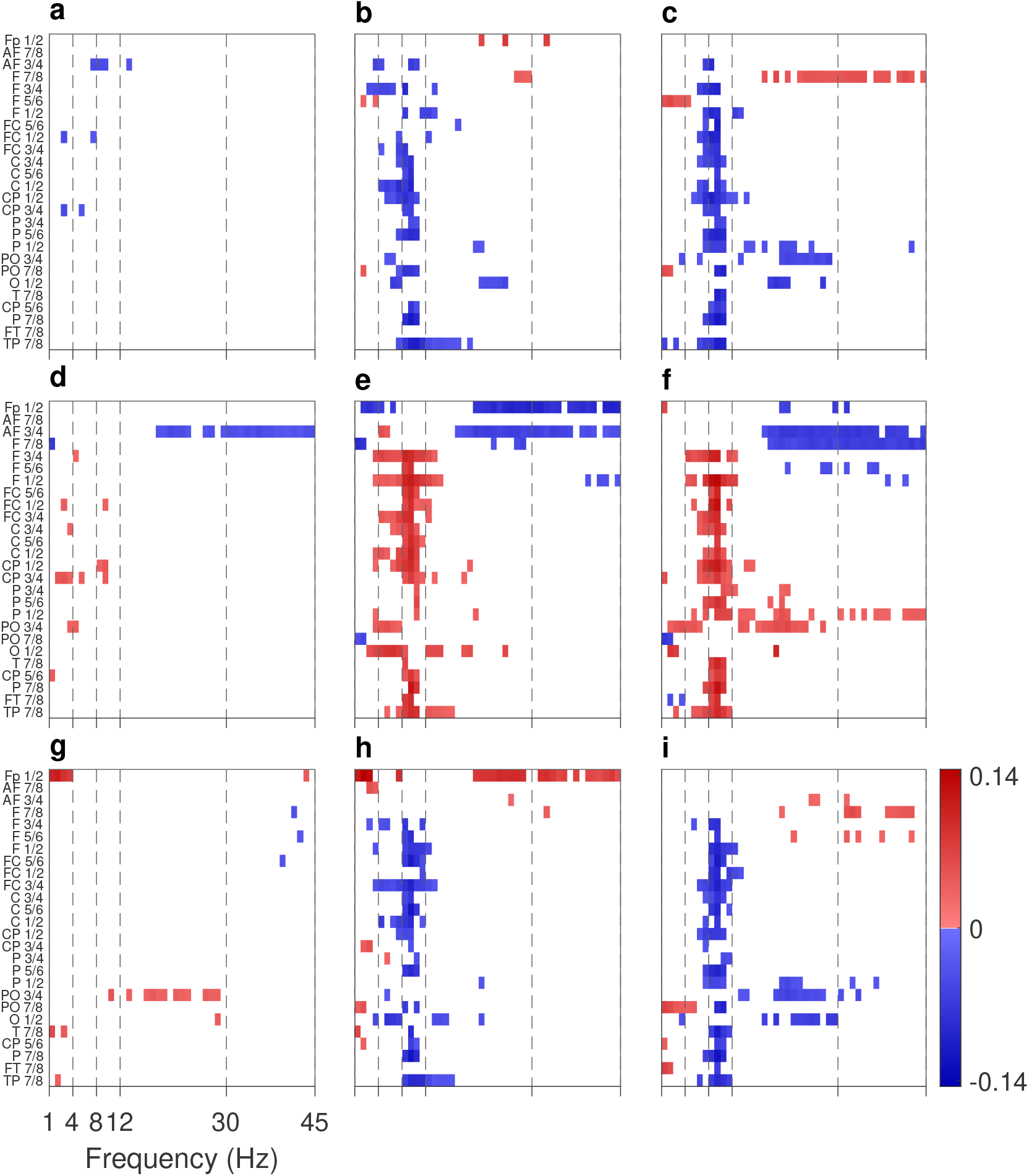
Plots of ΔMSC between symmetric electrode pairs versus frequency in control subjects. **(a)** MED-EO2_C_; **b** MED-EC1_C_; **c** MED-EC2_C_; **d** EO2_C_-EO1_C_; **e** EC1_C_-EO1_C_; **f** EC2_C_-EO1_C_; **g** EC2_C_-EC1_C_; **h** EO2_C_-EC1_C_; **i** EO2_C_-EC2_C_. All the other notations, etc. are exactly the same as in Fig. 2.

**Figure 10:**
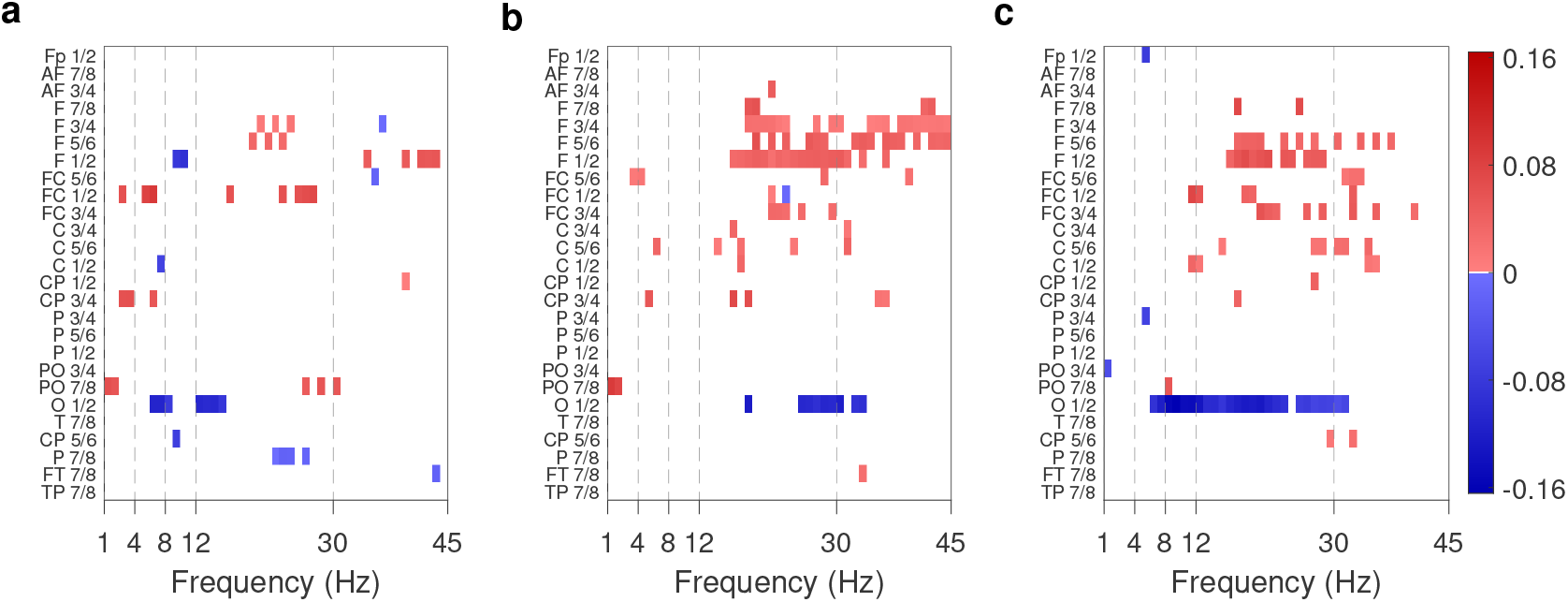
Plots of differences of MSC between meditators and control subjects versus frequency during **a)** initial eyes closed conditions (EC1); **b)** eyes closed conditions post MED and MUS sessions (EC2); and **c)** eyes-open conditions post MED and MUS session (EO2). All the figures are plotted with the same color scale. From the top to the bottom, the electrode pairs are arranged according to the anterior to posterior regions of the brain, with the temporal electrodes at the bottom. Only the electrode pairs and frequencies statistically significant with p ≤ 0.05 are highlighted with the color red (blue) indicating the increased (decreased) MSC in meditators than controls and others are shown as zero with white color in the figure.

**Figure 11:**
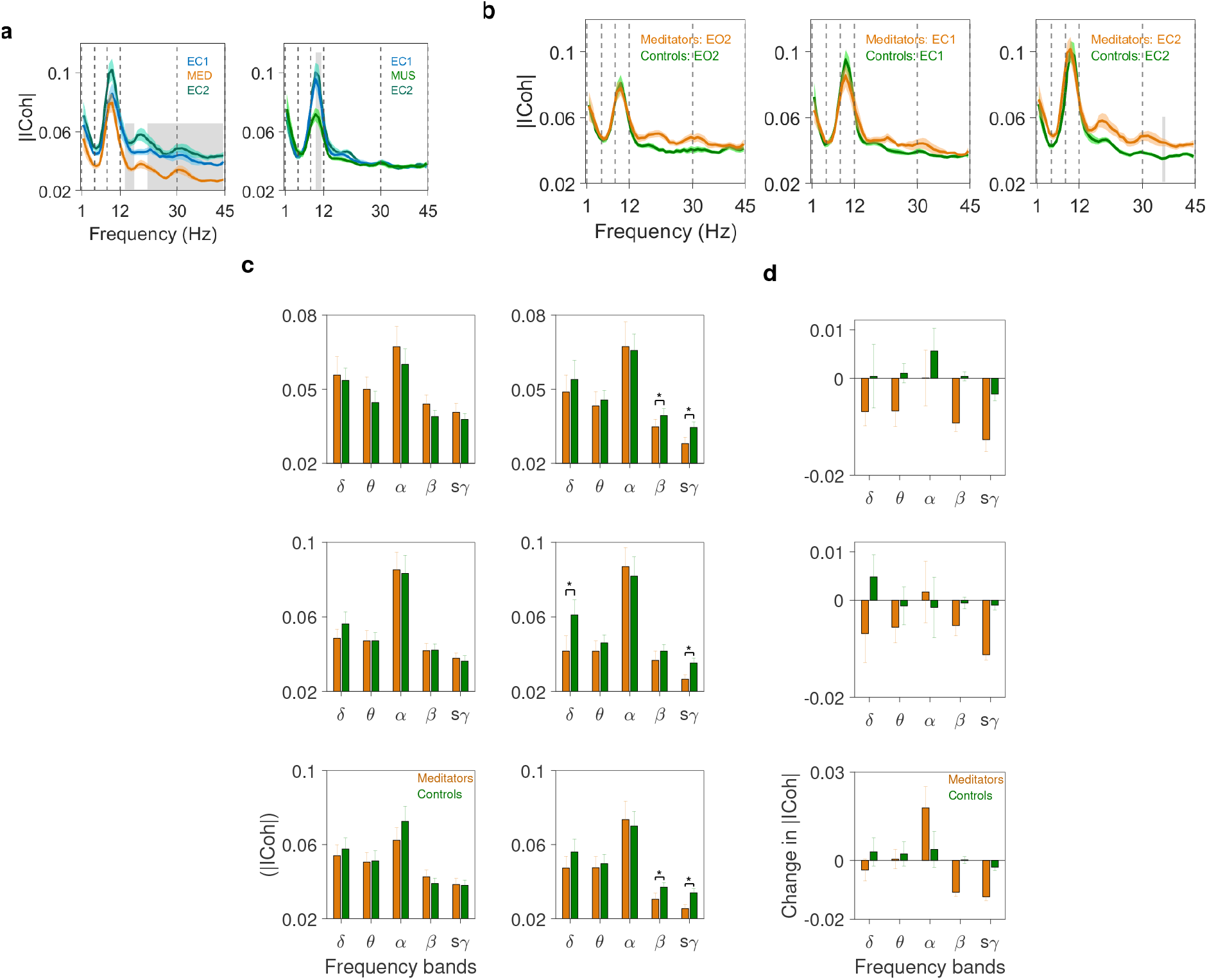
|ICoh| in meditators and controls: **a)** Grand averages of |ICoh| across all the electrode combinations as a function of frequency during different conditions within the groups. The respective light color band around each solid line represents SEM. Left column: meditators and right column: controls. **b)** Same as in **(a)** but across groups during EO2 (left), EC1 (middle) and EC2 (right) sessions. The vertical dashed lines indicate the boundaries of delta, theta, alpha, beta, and slow gamma (s*γ*) bands. The grey shaded area indicates the region of statistical differences between the conditions (EC1&MED, EC1&MUS) and groups after Bonferroni correction for multiple comparisons. **c)** Mean |ICoh| in frontocentral (top), centroparietal (middle) and temporal (bottom) regions during EO1 (left column) and during meditation or music sessions (right column) in different frequency bands. Error bars represent SEM. * represent the statistical significance after Bonferroni correction. **d)**: Change in |ICoh| in frontocentral (top), centroparietal (middle), and temporal (bottom) regions during meditation or music sessions compared to respective EO1 (MED-EO1_M_ and MUS-EO1_C_) in different frequency bands.

**Figure 12:**
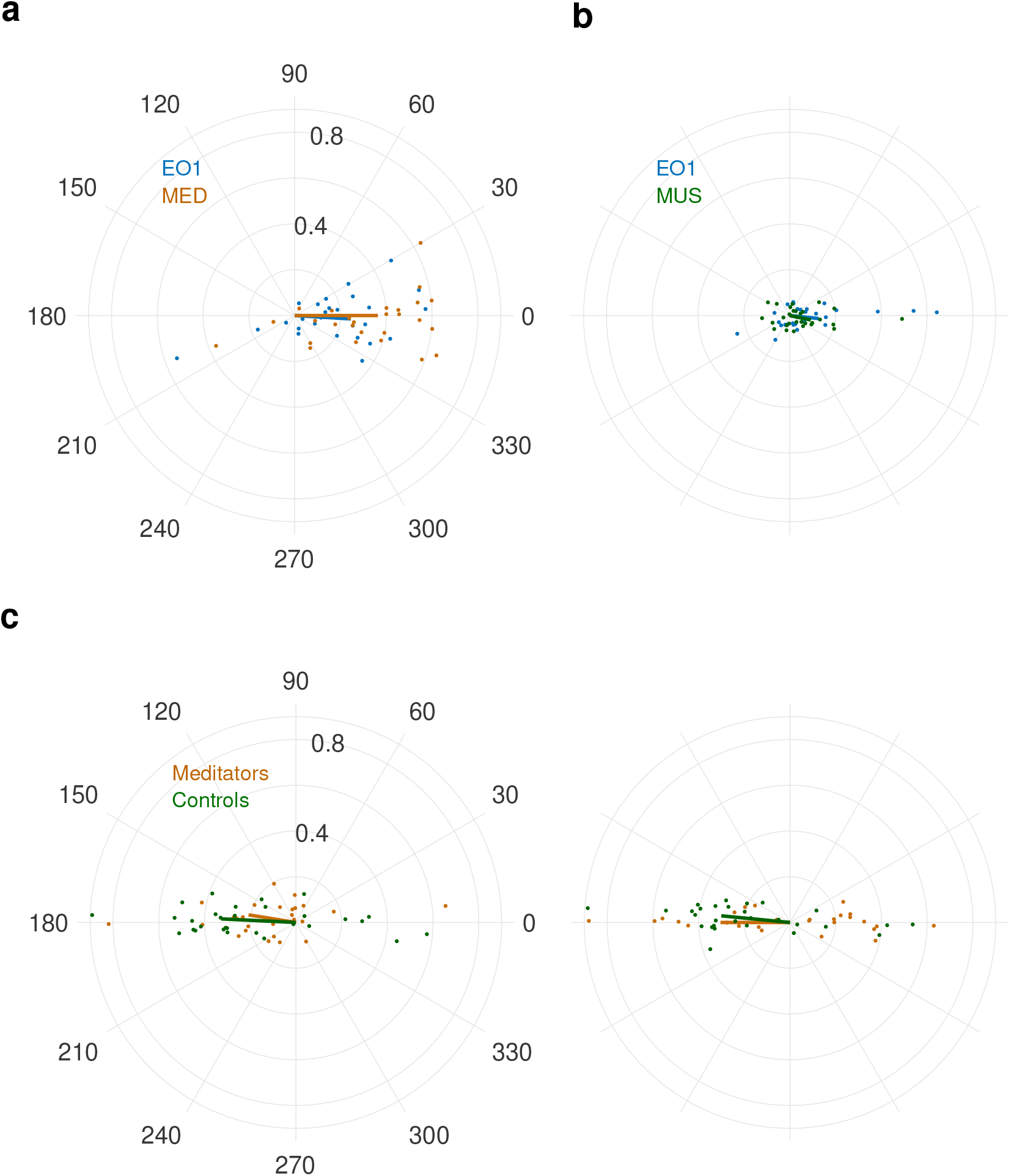
Polar scatter and polar plot presenting the relation between MSC and |ICoh| in meditators and controls. **a)** Scatter plots of points (meditators) in polar form of the coherency values of electrode pair FC5-FC6 at 9 Hz where MSC is higher during meditation than EO1. **b)** Same as in **a** but (controls) of electrode pair AF3-AF4 at 18 Hz where MSC is lower during music session than EO1. **c)** Scatter plots of points in the polar form of meditators and controls during EO1 condition (left) and during meditation/music sessions (right) in occipital electrode pair O1-O2 at 44 Hz. Solid lines in the respective color indicate the mean magnitude and phase.

**Figure 13:**
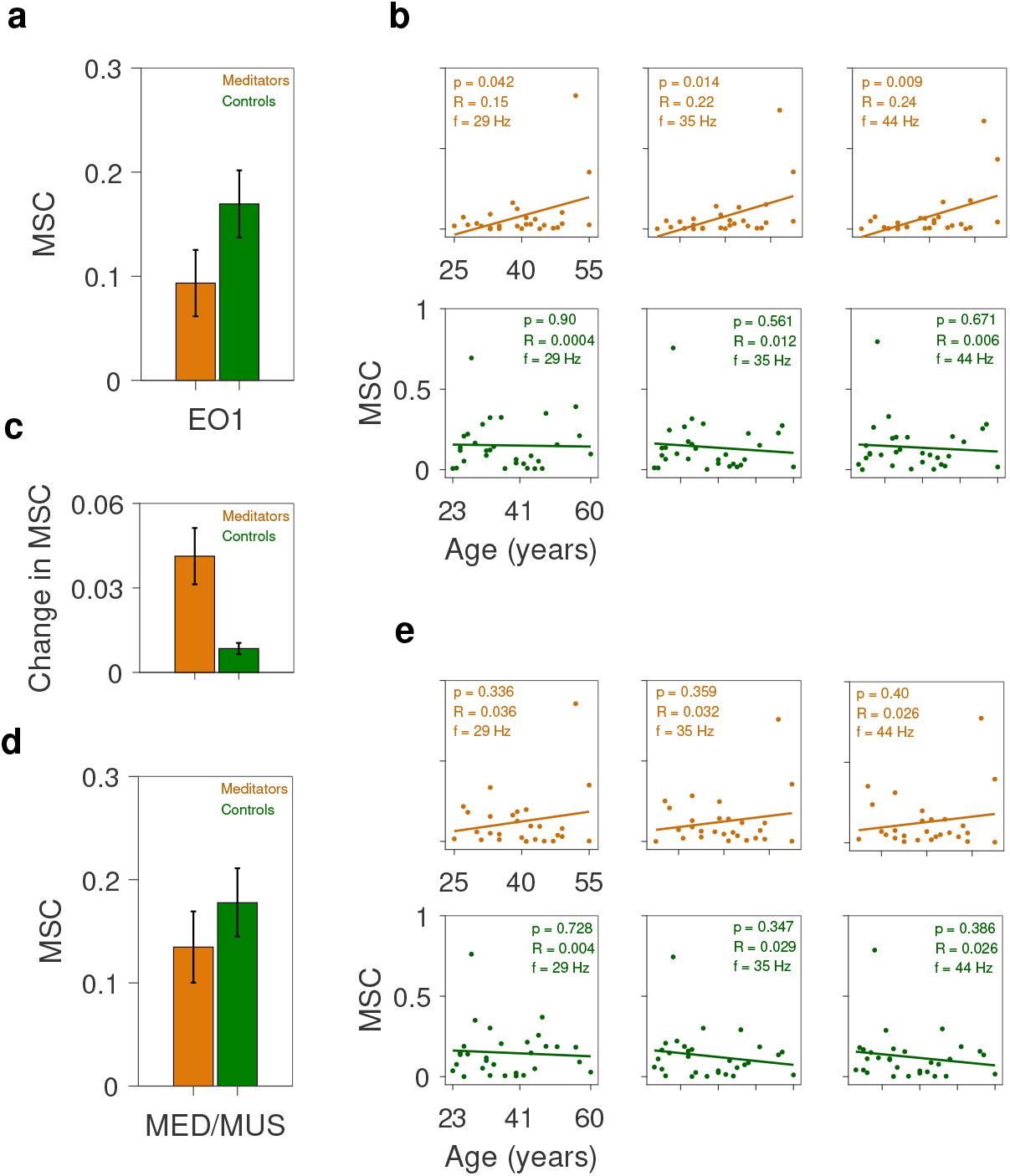
MSC between the occipital O1-O2 electrodes in meditators and controls: **a)** Mean MSC across all the bands (1-45 Hz) during baseline eyes-open condition. Error bars indicate SEM. **b)** Scatter plots of MSC during baseline EO condition as a function of the age of meditators (top row) and controls (bottom row) at three different frequencies. Each of the points represents a subject. Solid lines indicate regression fits, whose p-values are indicated in respective colors. (linear regression for MSC in meditators vs age: slope, *β*_1_ = 0.008 for 29 Hz; *β*_1_ = 0.008 for 35 Hz; *β*_1_ = 0.009 for 44 Hz. In controls, the slopes are always negative, *β*_1_ = -0.0003 for 29 Hz; *β*_1_ = -0.002 for 35 Hz; *β*_1_ = -0.001 for 44 Hz). **c)** Change in MSC with respect to the baseline eyes-open condition during meditation or music session. **d)** Same as in **(a)** but during meditation or music session. **e)** Same as in **(b)** but during meditation or music session. (In meditators, *β*_1_ = 0.004 for 29 Hz; *β*_1_ = 0.034 for 35 Hz; *β*_1_ = 0.003 for 44 Hz. In controls, *β*_1_ = -0.001 for 29 Hz; *β*_1_ = -0.002 for 35 Hz; *β*_1_ = -0.002 for 44 Hz).

### Changes in IHC in meditators compared with controls

Fig. 4 highlights the electrode pairs and frequencies with significant differences between the mean values of MSC of the two cohorts in initial eyes open condition (EO1) and during MED or MUS sessions. Fig. 4a shows that MSC values are higher in meditators in centroparietal region of the brain in higher theta, alpha, and low beta bands and beta band in frontocentral region. Lower mean MSC is seen in meditators than controls in occipital electrode pair O1-O2 which is consistent in EO1 condition in all the frequencies except delta and alpha bands. Fig. 4b shows the electrode pairs with significant differences in MSC between meditators and controls during the task (meditation and music segments, respectively). Significant differences are found in frontocentral and centro-parietal regions of the brain in the alpha band and in the frontal regions in beta and slow gamma bands. Only the electrode pair O1-O2 is found with higher MSC in controls which is statistically significant in the lower beta band. However, the increased MSC during the meditation session in meditators leads to no significant differences between the meditators and control subjects in the occipital region except in the lower beta band. This is because IHC was higher in controls in the higher frequencies during EO1, and there is no change during the task. The MSC in meditators was higher than controls in the initial eyes closed (EC1) condition in frontal, frontocentral region in the beta band. The MSC in O1-O2 pair was less in meditators in high theta and low beta regions as given in Suppelementary Fig. 10a. The decrease in MSC continues in O1-O2 pair during the post-meditation or music segment eyes open (EO2) condition (Supplementary Fig. 10c). In the post-meditation and music segments, MSC increases further in meditators in the frontal region as compared to controls in both EC2 and EO2 conditions in beta and slow gamma bands as seen in Figs. 10b and c. The occipital region MSC is higher in controls than meditators during all the baseline conditions at specific frequency bands.

The grand average of |ICoh| comparing the meditators and controls given in the left column of Fig. 5b suggests that there exists a difference in the EO1 condition across groups prominently in delta, beta and slow gamma bands. However, the differences are not significant at the Bonferroni significance level. The differences between the groups during task conditions i.e, MED and MUS session increases further as shown by the right column of Fig. 5b. It is observed that |ICoh| decreases in delta, theta, beta, and slow gamma bands but increases in alpha band. However, the significance at Bonferroni significance level of p ≤ 0.05/45 = 0.001 is found only at 16 Hz, 25-27 Hz, and 35-45 Hz. To further quantify the results of |ICoh| across groups, the |ICoh| values are averaged across different lobes; the results of frontal lobe (with Fp1-Fp2, AF3-AF4, AF7-AF8, F1-F2, F3-F4, F5-F6, F7-F8 electrode pairs) for EO1 and MED/MUS sessions are given in the top row, left and right columns, respectively, of Fig. 5c. The |ICoh| values are higher in meditators than controls in EO1 condition across all the bands except the delta. However, they are significant only in delta (p = 0.02) without Bonferroni correction and beta (p = 0.002) bands with Bonferroni correction. During MED/MUS session, the mean |ICoh| values are significantly different between the controls and meditators with Bonferroni significant levels of p ≤ 0.05/5 = 0.01 in slow gamma (p = 7.67*10^*−*4^), delta (p = 0.0017) and without Bonferroni significant levels in beta (p = 0.032). It is observed that |ICoh| decreases for meditators in all the bands and remains almost the same for controls except for the alpha band. Hence, except in alpha band, the mean |ICoh| values are more in controls than meditators. Significant differences were found in beta and slow gamma bands during EO2, EC1, and EC2 conditions across groups (Supplementary Fig. 11b).

We have further checked the changes in |ICoh| values across conditions in each group. The changes in |ICoh| values in frontal and occipital lobes during MED/MUS session compared with EO1 are given in Fig. 5d in the respective rows. During MED, |ICoh| in frontal lobe decreases significantly with Bonferroni significant levels of p ≤ 0.05/5 = 0.01 in delta (p = 0.004), beta (p = 3.23*10^*−*5^) and slow gamma (p = 1.09*10^*−*4^) bands, and without Bonferroni significant levels in theta (p = 0.04). In controls, the changes in frontal lobe |ICoh| values between EO1 and MUS session are not significant as shown in Fig. 5d (top row). The error bars represent the SEM. It is observed that the decrease in the |ICoh| in frontal lobe (top row) is higher in meditators than controls except in alpha band and is significantly higher in the beta and slow gamma bands. Alpha band change is the least in meditators and is the highest in controls. The mean |ICoh| during EO1 and MED/MUS in frontocentral, centroparietal, and temporal regions are given in Supplementary Fig. 11c and changes in |ICoh| during MED/MUS session compared to respective EO1 (MED-EO1_M_ and MUS-EO1_C_) is given Supplementary Fig. 11d.

### Reduced coherence in occipital electrodes in meditators than controls

The mean |ICoh| values calculated exclusively over three occipital electrode pairs PO3-PO4, PO7-PO8 and O1-O2 are given in Fig. 5c (bottom row) during the baseline EO1 (left column) and during MED/MUS conditions (right column). The |ICoh| values are not significantly different in any band between meditators and controls in EO1 condition. During MED and MUS sessions in meditators and controls respectively, the mean |ICoh| values of the occipital region showed significant differences between the controls and meditators with Bonferroni significant levels of p ≤ 0.05/5 = 0.01 in delta (p = 4.0*10*e*^*−*4^) and slow gamma (p = 1.47*10^*−*4^) bands, and without Bonferroni significant levels in theta (p = 0.038) and beta (p = 0.032). We have further checked the changes in |ICoh| values across conditions in each group. During MED, |ICoh| decreases significantly with Bonferroni significant levels of p ≤ 0.05/5 = 0.01 in delta (p = 9.14*10*e*^*−*4^) and slow gamma (p = 9.94*10^*−*5^) bands, and without Bonferroni significant levels in theta (p = 0.03) and beta (p = 0.04) bands. In controls, the changes in |ICoh| values between the EO1 and MUS session given by Fig. 5d (bottom row) are not significant.

Figure 6 compares the mean MSC values of occipital electrode pair O1-O2 across different frequency bands. In baseline conditions, the mean MSC of meditators is consistently less than that of controls across all the frequency bands (Fig. 6a)(left), but it increases during MED (Fig. 6b). The actual mean MSC during MED/MUS session is given in Fig. 6a(right). The increase in MSC during MED is significantly higher than that of controls occurring during the MUS session (Fig. 6b). The increase is mostly in lower and higher frequency bands except for the alpha and low beta band (Fig. 4b Fig. 6b).

To quantify the differences in O1-O2 electrode pairs, we have further tested by regressing MSC against the age of meditators and controls in higher frequency bands at three frequencies 29, 35, and 44 Hz both in baseline eyes open condition (Fig. 4a, Supplementary Fig. 13b) where MSC in meditators is significantly lower than controls and during meditation and music sessions (Fig. 4b, Supplementary Fig. 13e). When we pool the data across both genders, the results are significant and are moderately correlated (linear regression p ≤ 0.05) for meditators and poorly correlated in controls (Supplementary Fig. 13b). During MED, (Supplementary Fig. 13e) the correlation of MSC with age decreases and does not reach significance but the slopes remain positive at all the three frequencies calculated. In controls, the MSC is slightly more negatively correlated in the MUS session compared to EO1 condition. MSC measures at the above mentioned frequencies (29, 35, and 44 Hz) are also included in linear regression models to test whether a linear dependency exists with the meditation expertise. None of the regression models reveal a significant relationship between the MSC values and meditation expertise.

## Discussion

Coherence may be considered as a tool to determine the degree of synchronization which is the basis for information processing by the neurons (Von Der Malsburg, 1985). Measures of interhemispheric EEG coherence during a defined task reflect the FC of the hemispheres. It is known that different frequencies are dominant during different cognitive states of the brain like normal wakefulness, sleep, altered states of consciousness (eg. coma), and modified states of consciousness (eg. meditation, hypnosis). In this study, we have explored magnitude squared coherence (MSC) and the imaginary part of coherency (ICoh) between symmetric electrodes to study the cross-spectral correlation and information processing across the hemispheres. We have employed these two connectivity measures in order to investigate (1) whether there exists a baseline difference (trait effect) in interhemispheric connectivity, (2) the changes in the measures during the respective task conditions (meditation or music listening), (3) whether meditation and music listening sessions share some similarities while exhibiting distinct signatures. We have utilized complementary analytic approaches, which assess the MSC and ICoh of the EEG coherency, along with two within-subjects variables: frequency band (delta, theta, alpha, beta, and slow gamma), and 26 interhemispheric electrode pairs. In the present study, the absolute value of ICoh is used to convert the range to 0 to 1, similar to MSC.

### Differences in FC patterns of MSC and |ICoh| measures

The MSC and |ICoh| measures do not exhibit a similar pattern of changes since the former depends on both the amplitude and phase and the latter depends on the phase difference between the signals. MSC depends on phase consistency as well as amplitude correlations between the two signals. It is also possible that two signals are completely coherent but out of phase. The |ICoh| measure is the imaginary part of coherency given in equations (7) and (8). The phase difference between two signals plays a major role in estimating |ICoh| even when the magnitude is the same. The opposite trends of MSC and |ICoh| changes can be well understood from the supplementary Fig. 12. The polar scatter plots show the coherency values during different conditions (supplementary Figs. 12a and 12b) and across groups (supplementary Fig. 12c). However, the effect of volume conduction on MSC is reduced by applying the surface laplacian filtering (Perrin et al., 1989), which is considered to be one of the efficient methods to reduce the volume conduction effect in FC analysis. A study conducted to test the reproducibility of FC in resting-state EEG by Duan et. al (Duan et al., 2021) showed very low correlation between the MSC and |ICoh| with the definition of insensitive/sensitive to volume conduction. Hence the results obtained using MSC and |ICoh| may be discussed independently.

### Correspondence with previous findings

The increase in interhemispheric MSC across different regions in different bands during meditation may be due to the high emotional stability leading to implications on executive efficiency (Ganpat et al., 2011). This has also been attributed to increased brain idling and positive emotional experience during meditation. The increase in delta MSC which is locally distributed in pre-frontal, frontal, and frontocentral regions as compared to global distribution during sleep may be attributed to attention directed towards internal processing during meditation (Travis, 2014)(Horan, 2009)(Jaušovec, 2000)(Shaw, 1996). Studies also suggest the delta band engagement of interhemispheric connectivity between neural assemblies during the mental task involving complex calculations (Harmony et al., 1996). An increase in delta MSC during meditation also suggests its role associated with facilitating the mechanisms of large-scale integration during the meditative state (Lutz et al., 2008) with enhanced meta-cognition and vigilance resulting in increased access to neural networks involved in consciousness process (Raffone and Srinivasan, 2009)(Varela et al., 2001). The increase in FC in lower alpha band is in support of better functional network integration in experience meditators as measured with graph theory metrics (van Lutterveld et al., 2017). The results of the present study are in-line with (Murata et al., 2004) with respect to the significant increase in frontal interhemispheric EEG MSC in higher theta and lower alpha bands during meditation (Fig. 3a).

Enhanced MSC in frontocentral electrodes across hemispheres corresponding to supplementary motor area indicates information exchange due to visualized movements (an integral part of the Rajyoga meditation) considered to be a special form of motor behavior. Previous investigations have suggested that coherence in alpha band is a more sensitive parameter than amplitude or power value reflecting the meditative states (Dillbeck and Bronson, 1981)(Travis and Wallace, 1999). The increase in frontal EEG interhemispheric coherence in slow alpha has also been reported during Zen (Murata et al., 2004) and transcendental meditation (TM) (Orme-Johnson and Haynes, 1981), suggesting a common signature of higher internalized attention during meditation across various meditation techniques. Our study results are similar to these previous reports indicating that meditation modifies alpha EEG patterns significantly more than simple relaxation techniques. Significant reductions in alpha band phase lag between pairs of electrodes resulting in higher synchronization were found during TM in frontal and parieto-occipital areas compared to the resting eyes-closed condition proposing a phase synchrony model of consciousness and hypothesizing improved functional integration in information transfer involved in high-level cognitive processes (Hebert et al., 2005). (Stephan et al., 1995). Increased alpha coherence indicates higher functional coordination or information exchange/transfer (Petsche et al., 1988) between the brain regions and functional coupling (Thatcher et al., 1986). Slow alpha synchronization is related to the inner-directed, internalized attention during meditation and high-alpha desynchronization is related to external attention which may not be involved in meditation (Shaw, 1996). Enhanced MSC during meditation in higher theta and alpha bands in prefrontal and anterior frontal electrodes corresponding to the prefrontal cortex may be related to the emotional component as suggested by (Davidson, 2004) and also the attribution of intention to others (Brunet et al., 2000).

It is important to note the different results directions of the two measures MSC and |ICoh| and establishing the independence between the two measures requires further investigations by considering all possible electrode pairwise combinations and cluster-wise analysis. While there is no parallel between the previous meditation studies on Rajyoga practice and the present findings, increased |ICoh| measure was reported across three meditation techniques in delta networks (Yordanova et al., 2020). Different meditation traditions with EEG measurements are known to present different characteristics. However, in the present study, a significant decrease is observed in beta and gamma bands which are in line with a study on five different meditation traditions using lagged coherence which has close mathematical relation with ICoh, reported decreased FC during meditation (Lehmann et al., 2012). The decrease in ICoh connectivity may imply the subjective experiences comprising functions such as autobiographic memory, out-of-body experience (OBE), and self-awareness (Conway, 2005),(Blanke and Arzy, 2005). The ICoh connectivity was not significantly changed during the OBE under hypnosis but a decrease in power was observed in beta and gamma bands (Facco et al., 2019). Future work will have to examine possible specifics of these changes as Rajyoga meditation follows the concept of the out-of-body process for further concentration and realization during meditation.

Interhemispheric alpha MSC was found to be inversely correlated with the psychomotor poverty scores in a study conducted on 37 neuroleptic-naive patients with recent onset of illness (John et al., 2002). The inverse correlation was found in central and occipital regions in the eyes open condition indicating asymmetric cortical hypofunction associated with symptoms of schizophrenia. Further, studies that had demonstrated negative correlation between negative symptoms and interhemispheric alpha coherence reported a reversal of these findings following drug treatment (Merrin and Floyd, 1992). The increase in MSC across all the bands except alpha during meditation in the occipital electrodes O1-O2 may be attributed to the enhanced visual-spatial processing and recognition (Nagendra et al., 2015). The non-increase in alpha band may be attributed to the unique practice of meditation with eyes open condition which needs further investigation to establish the nuances between Rajyoga and other meditation practices during the baseline eyes open and closed conditions. The change in controls during the music listening session compared to the eyes open baseline conditions are confined to the decreased MSC in anterior-frontal electrode pair AF3-AF4 in high beta and slow gamma bands (Fig. 3b). However, we are unable to relate the results of the present study with any of the previous studies on MSC during music listening.

Distinct connectivity patterns are observed in meditators and controls during baseline eyes-open and meditation or music sessions. The differences are significant in higher bands mostly in fronto-central and centroparietal regions. Significant theta MSC increase in central and centrotemporal regions during meditation compared to rest condition was reported in (Lagopoulos et al., 2009). The MSC between occipital electrodes O1-O2 is significantly lower in meditators than controls in baseline eyes-open condition. This may be a trait effect in Rajyoga meditators due to the practice of meditation with their eyes open. However, this requires further study to analyze the altered visual sense and information processing in Rajyoga meditators. MSC in occipital electrode pair is partially positively correlated with the age of meditators, but not with the meditation experience. This may again be a trait change due to the practice of meditation with eyes open. The connectivity patterns during the task conditions are different in the two groups. Meditators have distributed increased MSC in alpha band and decreased MSC in slow beta band in the occipital region. The results of our study based on the synchronous activity of EEG between the two sides of the brain extend the previous studies and suggest that the brain networks related to symmetric electrodes are different in meditators from the age-matched controls indicating the trait effect.

### Possible underlying mechanisms and relevance

If we assume that the observed increased or decreased MSC in (most of) the meditators across different regions of the brain is caused due to long-term practice, the underlying mechanism needs to be answered. A study conducted to investigate the role of alpha waves in sensory processing reported that alpha oscillations in the frontal lobe might be involved in the origin of top-down control thereby regulating the perceptual gains which may be related to the increased processing of different sensory information during meditation (Misselhorn et al., 2019). A groundbreaking study in cognitive neuroscience has been on the changes in alpha oscillations in the visual regions of the brain when humans close their eyes (Berger, 1929)(Adrian and Matthews, 1934) and may be related to the ongoing visual processes. It is understood that the decrease in alpha oscillations when humans open their eyes is due to the excitation of visual sensory information and alpha oscillations are considered to be the representation of the no-input condition through inhibition (Pfurtscheller et al., 1996). The decreased MSC between symmetric electrodes O1-O2 in eyes open condition of long-term meditators compared to controls may be related to the efficient visual information processing. The increased MSC during meditation may indicate effective gating of the visual information even with the eyes open. However, the increase in MSC except in alpha band may reflect the consequence of internal visualization involved in Rajyoga meditation practice. Further analysis shall be carried out in this direction to decode the neural mechanisms.

Similar observations indicating the relationship between the age-related decrease in controls but not in long-term meditators have been previously reported with respect to different connectivity measures. It is reasonable to accept that some brain regions or FC or networks in long-term meditators are changed through the training whereas in matched healthy controls, they are maintained as they are. Nevertheless, it is worth noting that the group differences in baseline conditions are confined to specific brain regions including the frontal in alpha oscillations and occipital lobes in oscillations other than the alpha band. Thus meditation appears to be an effective mental practice with the potential to change the FC of the brain at large. Collective evidence suggests that regular meditation as an intervention may be beneficial to people suffering from different psychological disorders. However, given the sparsity of the data, meditation studies (especially longitudinal studies) in healthy populations may give more evidence for considering meditation in clinical intervention studies.

### Limitations and implications for future research

This study has the following limitations in methodology which affect the analysis on estimating the MSC and |ICoh| measures. (1) The baseline eyes open and eyes closed conditions before and after meditation or music segments are not randomized. (2) The same music is played for all the control subjects, while the preference of subjects may differ. (3) The duration of baseline and task conditions are not matched because of the practical issues of making a subject sit idle and relaxed during baseline conditions. (4) The meditation and music session duration are not matched. The control subjects were unable to rest or relax without getting to sleep when the music session was long and hence the duration was reduced from 10 to 5 minutes. (5) Since MSC and |ICoh| measures depend on the number of trials or epochs during a condition, the analysis epochs of the meditation condition have been reduced to 5 minutes. Future work shall include testing the patterns in executive control networks such as frontoparietal networks and medial frontal networks which involve cognitive control, and attention regulation, and reporting the results on intra-hemispheric and anterior to posterior FC analysis.

## Materials and methods

### Subjects and EEG recordings

The dataset consists of 27 (11 female) healthy right-handed meditators (mean age: 39.8 years SD: 8.4 years) with an estimated lifetime meditation experience of 14.2 ± 9.2 years. They were recruited from different meditation centres in Bangalore through the centre trainers. We also recruited 30 (12 female) healthy right-handed controls (mean age: 37.3 years SD: 10.5 years) from the student and staff community of the Indian Institute of Science. Participants who had no previous experience of any type of meditation were recruited through awareness talks on meditation studies. Both meditators and controls were screened for any history of general health issues, and neurological/psychiatric illness. All the subjects had normal or corrected to normal vision and participated in the study voluntarily. Informed consent was obtained from the participants and they were briefed about the study before performing the experiment. The ages of meditators and controls matched with p = 0.243 tested with the Wilcoxon rank-sum test. Ethical clearance for the experimental protocol was obtained from the IISc Institutional Human Ethics Committee, with IHEC number 02/20201126.

The raw EEG data was recorded using a 64-channel waveguard cap and ANT Neuro mylab acquisition system. Electrodes were placed according to the international 10-10 system. The data were recorded with a sampling rate of 1000 Hz. Although the impedance threshold in the acquisition system was set to 20 kΩ, the electrode impedance was maintained at less than 10 kΩ during recording. EEG signals were referenced to CPz during acquisition. Other physiological signals, namely ECG, respiration and GSR were also recorded with the same acquisition system.

### Experimental setup

The participants sat in a dark room with the comfortable seating of their choice. None of them sat in a cross-legged position. Meditators and controls performed the instructed task as shown in Fig. 1a top row and bottom row respectively, which was a within-subject design. Subjects were given enough time to adapt to the conditions of the recording room while the experimenters briefed them about the experiment. The average preparation time for the recording was approximately 30 minutes. Subjects were instructed to minimize movement during the recording and baseline recording was performed for 5 minutes each with eyes open (EO1) and closed (EC1) conditions before the meditation or music session. Meditators practiced BKRY meditation (MED) for approximately 10 minutes, and control subjects listened to music (MUS) as a comparable task. The music was selected from the original soundtrack of the playlist on youtube channel of the Brahmakumaris organization and is also being used by a few meditators of this school of meditation as background music to aid the practice in the initial years. Music listening was considered as the task comparable to meditation in the present study since many intervention studies have reported the effect of music listening as a good task to compare with meditation (Henneghan et al., 2020),(Nagamatsu and Ford, 2019),(Innes et al., 2016). Final baselines were again recorded with eyes closed (EC2) and open (EO2) conditions for a duration of 5 minutes each. The eyes open and closed segments were not randomized since the meditation and music segments were recorded with the eyes open condition.

### Rajyoga meditation

BKRY meditation is taught by the Prajapita Brahmakumari Ishwariya Vishwa Vidyalaya that has consultative status with UNICEF, UNO, and WHO (Kiran et al., 2005). It is an open-eyed meditation technique that involves focusing on oneself as a peaceful being of light at the center of the forehead (Ramsay et al., 2010), (Brahmakumaris, 2022). By this practice, the cluttered mind becomes calm and peace is experienced (Sharma et al., 2018a), (Telles and Desiraju, 1993). As one advance in this practice, the ability to perceive as a detached observer is mastered. Any object or circumstance including the body, breath, thought, emotion, and situation can be managed efficiently as a detached observer. This state of being a detached observer is called Sakshi bhav. BKRY meditation has been practiced with various stages (University, 1994) namely relaxation, concentration, contemplation, realization, and meditation (Ramesh et al., 2013) (appendix of (Ganesan et al., 2020)), (Patel, 1984). The time required to attain a given state reduces with experience. Among many masteries one gains through Rajyoga, sakshi bhav is one of the key skills that has many practical implications in daily life. It is an induced modified state of consciousness with heightened self-awareness, alertness, and control of thoughts. As the individual practices with eyes kept open, the external visual stimuli are altered with the advance in the meditation practice session. This may lead to reduced information processing of external stimuli involving sensory gating. Memory access may be modified occasionally and could be normal during the mind-wandering segments. An audio recording prepared by a senior BK teacher was provided to all the meditators and they were asked to practice at least two times before the recording session in order to keep the practice variables and the duration of the steps aligned. No audio recording was played during the recording session for meditators.

### Data conditioning and artifact rejection

The MILE (Meditation in Lab Environment) preprocessing pipeline is illustrated in Fig. 1b. Preprocessing was carried out using custom MATLAB (The MathWorks Inc., Ver 2020b) scripts based on EEGLAB version v2021.1 (Delorme and Makeig, 2004). The raw EEG data is highpass filtered at 1 Hz to remove the DC component and then notch filtered with F_center_ = 50 Hz with bandwidth = 2 Hz to remove line noise. Hamming windowed sinc filter is used for both high-pass and notch filtering. The filtered data is then epoched with segments of 500 ms duration with an overlap of 50%. Bad channels are detected as given in Supplementary Fig. 7a. Power spectral density (PSD) is using Fast Fourier transform (FFT) with multi-tapers using Chronux (Mitra, 2007) toolbox and averaged across epochs resulting in one PSD vector for each channel. The global mean is computed across channels and bad channels are detected when 30% of a channel’s frequency bins lay outside the mean ± 2 std. Bad segments are detected as given in Supplementary Fig. 7b. Three features are used for bad segment detection (1) Time Domain: The mean voltage per channel is calculated over all the epochs and epochs with voltage exceeding mean ± 6 std are marked for rejection; (2) Frequency Domain: The mean PSD for each channel is calculated over all the epochs and epochs with PSD exceeding mean ± 3 std are marked for rejection and (3) Statistical features: The waveform length and mean frequency are calculated over all the channels epochwise. A histogram is obtained for the features channelwise and an envelope is fit on the histogram bins. The envelope is used to detect outliers in the histogram. The bad segments obtained from the above three methods are combined using logical OR obtaining a bad-segment matrix of size n_channel_ * m_epochs_. Epochs with bad segments greater than 0.1*n_channel_ are finally labeled as bad segments.

Independent component analysis (ICA) is applied using runica algorithm ((Makeig et al., 1997)) after removing the bad channels and segments. The ICA components are then arranged according to variance and the top 20% of the ICA components are considered for rejection. The ICA components are then labeled using ICLabel (Pion-Tonachini et al., 2019), an automated independent component classifier toolkit resulting in a label (muscle, eye, brain, heart, channel noise, line noise, other) for each component. Component labels exceeding the thresholds (muscle - 0.7, eye -0.7, heart - 0.5) are marked for rejection. After removing the labeled ICA components, the data is low-pass filtered to 95 Hz. Further, the rejected bad channels are interpolated using the spherical interpolation function available in EEGLAB. Finally, the data is visually inspected to remove any artifact which may still be present. With all these conditions, the rejection was less than 10% of the data (4.2 ± 2.7% for meditators and 6.5 ± 4.5% for controls datasets). The data of all the subjects are derived using three distinct references - the average of mastoid electrodes M1 and M2, the common averaging, and reference electrode standardization technique (REST) (Dong et al., 2017) - to verify the effect of the choice of reference on the connectivity measure. The effect of volume conduction on the computation of EEG FC measures is minimized using spatial filtering. Surface Laplacian filtering was applied with Legendre polynomial of order 50 and smoothing parameter (*λ*) of 10^*−*5^ (Perrin et al., 1989).

### Data analysis

The primary goal is to analyze the IHC as an FC measure across conditions within and between the groups. In this study, we have used reliable methods which have a good mathematical basis. We have used sensor-level/channel space analyses with reduced volume conduction effects instead of source level for which the source localization techniques need to be applied. 26 electrode pairs (depicted in Fig. 1c) and frequencies from 1 to 45 Hz are considered for the IHC analysis. An interhemispheric electrode pair is discarded if any one of the electrodes is marked to be bad in more than 10% of the subjects in any group. We analyze the data using custom codes written in MATLAB and Chronux toolbox. We use non-overlapping epochs of 1000 ms duration with a frequency resolution of 1 Hz for the FC analysis. The frequency bands considered are delta (*δ*) [1-4] Hz, theta (*θ*) [5-8] Hz, alpha (*α*) [9-12] Hz, beta (*β*) [13-30] Hz and slow gamma (s*γ*) [31-45] Hz. Since the measures used in the study are sensitive to the number of epochs considered, the number of epochs of the meditation segment is matched to the number of epochs in initial eyes-open baseline conditions by considering the middle portion of the meditation segment. This approach is considered as opposed to the initial and final portion of the meditation segment by allowing sufficient time to elapse after the start of the meditation session. The number of epochs analyzed for each condition is listed in Table 1.

**Table 1:** Number of epochs (mean and standard deviation across the subjects) considered for the study after preprocessing.

| Condition | EO1 | EC1 | MED/MUS<br>available | MED/MUS<br>considered | EC2 | EO2 |
| --- | --- | --- | --- | --- | --- | --- |
| Meditators | 288±25 | 286±26 | 745±164 | 288±25 | 248±73 | 246±67 |
| Controls | 295±29 | 299±21 | 304±58 | 304±58 | 293±37 | 276±42 |

### Spectrum and cross-spectrum

For each electrode pair considered in Fig. 1c, the power spectral density *S*_*ii*_ and cross-spectral density *S*_*ij*_ are estimated using Fourier transform with multi-tapers using Chronux toolbox, at all the frequencies from 1 to 45 Hz, both included.

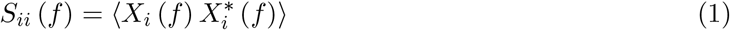

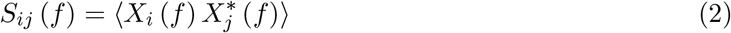

where *X*_*i*_ (*f*) and *X*_*j*_ (*f*) are the Fourier transform values at electrodes *i* and *j*, respectively, at frequency *f*, * denotes complex conjugation and ⟨·⟩ denotes the expectation over all the epochs in the state under consideration (Cohen, 2014).

### Magnitude squared coherence (MSC)

Coherency *C*_*ij*_ (*f*) between the electrodes *i* and *j* is given by

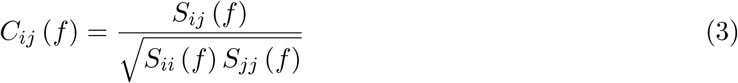

Using equations (1) and (2)

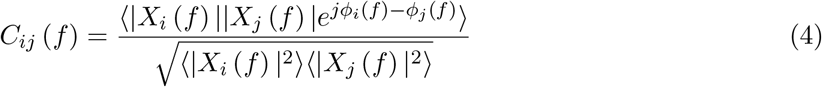

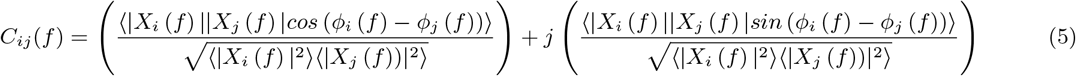

where |*X*_*i*_ (*f*) | and *ϕ*_*i*_ (*f*) are the magnitude and phase of *X*_*i*_ (*f*). Coherence *Coh*_*ij*_ (*f*), is defined as the absolute value of coherency and magnitude squared value of coherency (MSC) given by:

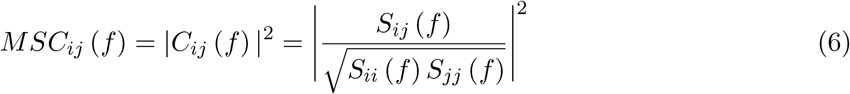

MSC ranges between 0 and 1, with a value of 1 indicating complete correlation and 0 indicating total absence of correlation. In this study, *i* and *j* are symmetric electrodes, i.e, one from the left hemisphere and the other from the symmetric location on the right.

### Imaginary part of coherency, ICoh

Proposed by (Nolte et al., 2004), ICoh is computed using only the imaginary part of the coherency given in equation (3) to study the interactions between different brain regions. ICoh is considered to exclude coherent sources with phase lag of zero thereby reducing the effect of field spread due to volume conduction. ICoh is defined as

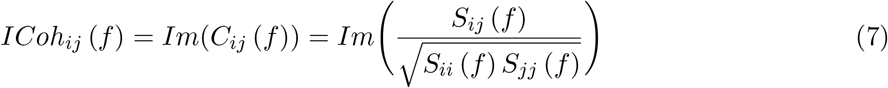

From equation (5)

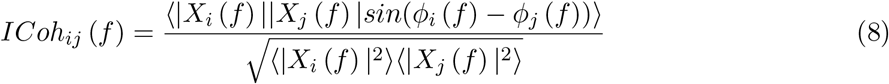

where *Im* stands for the imaginary part. The value of ICoh ranges from -1 to 1. A positive value implies that the electrodes *i* and *j* are interacting and *j* is earlier than *i*, indicating the flow of information is from electrode *j* to *i* and if negative, vice versa. Further, since the value of ICoh is proportional to the sine of the angle between the two channels at that frequency, it can be distinctly different from MSC, which is a magnitude measure. We use the absolute value of ICoh (|ICoh|) as a different measure of connectivity. After three-point smoothing, (|ICoh|) values of all the electrode pairs are pooled together to obtain the grand average (Yordanova et al., 2020) and also lobewise pooled to present more generalized data presentation by performing data reduction.

### Effect of referencing on MSC

Magnitude squared coherence (MSC) is a quantitative measure of the similarities between two signals in the frequency domain (Bortel and Sovka, 2006). MSC is affected by volume conduction and hence spatial filtering is performed to reduce its effect. Apart from the issue of field spread, coherence estimation may be erroneous due to reasons such as less number of epochs and effects of referencing. It is reported by (Nunez et al., 1997) that the average mastoid referencing adds from 0.1 to 0.3 to the coherence estimates, especially more for interhemispheric coherency estimated from electrode pairs close to the reference locations (mastoids). Figs. 2a, 2b, and 2c display the connectivity values for average mastoids, common average and REST referencing, respectively. REST referencing, proposed by (Dong et al., 2017), transforms the multi-channel data to approximately zero reference. It is evident from the figures that the coherence estimated in the peripheral electrode pairs near the mastoid electrodes, namely AF7-AF8, F7-F8, PO7-PO8, T7-T8, and FT7-FT8, are higher with the mastoid reference than the values estimated with other references. Hence, common averaging referencing was considered for both the FC measures in the present study.

MSC and (|ICoh|) are estimated for all the frequencies from 1 to 45 Hz with a resolution of 1 Hz. MSC values between the 26 pairs of electrodes for 45 frequency points for each subject result in 1170 values considered for the study. Only the statistically significant electrode pairs are highlighted in the results and the values of the non-significant pairs are made zero.

### Statistical significance

Non-parametric tests are used to evaluate the statistical significance of the results obtained since the values do not come from a normal distribution as determined using the Kolmogorov-Smirnov test in MATLAB and do not meet the parametric test requirements. To compare the different states (MED with EO1, EO2, etc.) of the same sample set, Wilcoxon signed-rank test (Wilcoxon, 1992) is used. To compare the results of different sample sets (namely controls versus meditators), Wilcoxon rank-sum test (Wilcoxon, 1992) is used which is equivalent to Mann-Whitney U-test. A result is considered to be statistically significant if p ≤ 0.05 for the 2-tailed hypothesis. Wherever necessary, false detection rate (FDR) correction is applied (Benjamini and Hochberg, 1995) to resolve the problem of multiple comparisons.

## Conclusion

Our experimental results show a clear change in the interhemispheric coherence pattern with two coherency-based FC measures during meditation indicating a possible change in the state of consciousness of the practitioners. In summary, interhemispheric MSC is higher and |ICoh| is lower in meditators than the mean age-matched control subjects implying that the effect of meditation is to preserve the EEG spectral synchrony. The results show that the increased MSC in the alpha band in frontocentral and centroparietal regions is associated with the activation of attention networks and may indicate functional integration. The increase in the MSC in alpha band in the central region in one electrode pair shows that meditation activates neural structures involved in motor movements suggesting the role of visualization. The results of this study suggest a new approach to utilizing interhemispheric coherence in meditation studies.

## Acknowledgements

The authors sincerely thank Rajayogini BK Ambika, Rajyogi BK Srikant, and the local centre teachers for facilitating and encouraging meditators to participate in the study. Authors also thank the control subjects and community members of IISc for volunteering in the study. Authors are grateful to Prajapita Brahmakumaris World Spiritual University, Mt. Abu, for lending their EEG system unconditionally for carrying out this study.

## Data and code availability statement

The EEG data used in the proposed study was recorded as a part of a postdoctoral fellowship that involved other questionnaires, experiments, and physiological signals such as ECG, GSR, and respiration, whose analyses are still in progress. Hence, the data used is not publicly available according to the rules and regulations of the fellowship. All the coherence analyses were performed using the Chronux toolbox (version v2.10), available at http://chronux.org.

